# Structural Basis for the Phase Separation of the Chromosome Passenger Complex

**DOI:** 10.1101/2023.05.22.541822

**Authors:** Nikaela W. Bryan, Aamir Ali, Ewa Niedzialkowska, Leland Mayne, P. Todd Stukenberg, Ben E. Black

## Abstract

The physical basis of phase separation is thought to consist of the same types of bonds that specify conventional macromolecular interactions yet is unsatisfyingly often referred to as ‘fuzzy’. Gaining clarity on the biogenesis of membraneless cellular compartments is one of the most demanding challenges in biology. Here, we focus on the chromosome passenger complex (CPC), that forms a chromatin body that regulates chromosome segregation in mitosis. Within the three regulatory subunits of the CPC implicated in phase separation — a heterotrimer of INCENP, Survivin, and Borealin — we identify the contact regions formed upon droplet formation using hydrogen/deuterium-exchange mass spectrometry (HXMS). These contact regions correspond to some of the interfaces seen between individual heterotrimers within the crystal lattice they form. A major contribution comes from specific electrostatic interactions that can be broken and reversed through initial and compensatory mutagenesis, respectively. Our findings reveal structural insight for interactions driving liquid-liquid demixing of the CPC. Moreover, we establish HXMS as an approach to define the structural basis for phase separation.

## Introduction

Membraneless intracellular compartmentalization is central to a long and growing list of biochemical transactions at diverse sub-cellular locations ^1^. Proteins are by-and-large the drivers of the formation of these compartments, but there is debate about whether their underpinnings are either low-affinity/low-specificity interactions yielding phase separation ^2^ or multivalent site-specific interactions ^3^. By its very nature, ascertaining the former type of interaction is essentially intractable by conventional structural biology methodologies. Moreover, the latter type of interaction is typically beyond current structural approaches, since multivalency involving contacts with highly flexible surfaces confounds traditional methodologies used in structural studies. Approaches that provide mechanistic insight on specific complexes engaged in membraneless compartmentalization are highly limited to date, especially when the molecules undergoing putative phase separation are more complex than a relatively small individual polypeptide. There is some reported success with crosslinking approaches ^4^ and NMR ^5–7^ but deciphering the physical basis of membraneless compartmentalization will ultimately require advancing new technologies and/or new applications of existing ones. The proposed types of bonds that are involved in phase separation are similar to those involved in conventional protein folding and interactions (e.g. hydrophobic and electrostatic interactions) ^1^; however, specifically how the structure and dynamics of a protein or protein complexes are impacted upon engaging in higher-order interactions is almost entirely unknown.

One proposed cellular compartment is the inner centromere, comprised, in part, of chromatin and the chromosome passenger complex (CPC) (Trivedi et al. 2019). The CPC is one of the key regulators of cell division and is comprised of four subunits: the serine/threonine enzymatic core Aurora B kinase, and three regulatory and targeting subunits, the scaffold inner centromere protein (INCENP), Survivin, and Borealin (also known as Dasra-B) ^9^. The activity of the CPC is strongly based on its sub-cellular localization during specific stages of cell division. In particular, during prometaphase the CPC is strongly localized to the chromatin spanning the two replicated centromeres, called the inner centromere. At the centromere, the CPC is involved in the process of mitotic error correction, whereby misattachments of centromeres to the microtubule-based mitotic spindle are rectified ^10^. Conventional targeting mechanisms through molecular recognition is required for CPC localization to the inner centromere. Specific chromatin marks at the inner centromere are recognized to direct CPC localization: the Survivin subunit directly binds H3^T3phos^ and the adaptor protein, Sgo2, indirectly binds to H2A^T120phos^ ^11–13^. The three non-catalytic subunits of the CPC (INCENP^1–58^, Borealin, and Survivin) form soluble heterotrimers that have a propensity to undergo liquid-liquid phase separation ^8^. Deletion of one region of Borealin between amino acids 139-160 (Borealin^Δ139–160^) or disrupting the strong positive charge in this region disrupts phase separation in vitro. These mutations within Borealin also reduce CPC accumulation at the inner centromere and its ability to robustly bundle spindle microtubules ^8, 14^. Furthermore, this region of Borealin overlaps with its mapped protein surface that contributes to nucleosome binding of the CPC ^15^. Besides the requirement for this region of Borealin, nothing mechanistic is known regarding how the CPC phase separates.

Here, we measure the change in polypeptide backbone dynamics of the INCENP/ Borealin/Survivin heterotrimer (ISB) in either a soluble or liquid-liquid demixed state using hydrogen/deuterium exchange coupled to mass spectrometry (HXMS). The most prominent changes in backbone dynamics are measured as additional protection from HX, primarily localized to discrete portions of α-helices of the INCENP and Borealin subunits. By combining information learned from peptide mapping provided by the HXMS data, a stepwise candidate mutagenesis approach, high-resolution structural information from the crystal packing behavior of ISB heterotrimers, and biochemical complementation, we identify three separate salt-bridges that drive liquid-liquid demixing.

## Results

### HXMS Identifies Regions with ISB Heterotrimers Impacted by Phase Separation

The ISB heterotrimer is comprised of the N-terminal 58 amino acids of INCENP, along with both full-length Survivin and Borealin (Fig. 1A). Together, it forms a three-helix bundle, containing a histone-binding module from the Survivin subunit and a C-terminal extension of Borealin that is reported to be mostly unstructured ^16^. Prior ISB phase separation was performed by either the addition of a polymeric crowding agent or by lowering the ionic strength ^8^. Polymers, like those typically used in phase separation studies (e.g., polyethylene glycol or dextran), are incompatible with the mass spectrometry step in HXMS that we intended to use to study the ISB, since the resulting spectra from polymers obscure those from the peptides under investigation. Thus, we studied the phase separation properties of the ISB over a range of protein concentrations and ionic strengths in the absence of polymeric crowding agent (Supplementary Fig. 1). From this, we focused our initial attention on a condition (25 μM ISB, 75 mM NaCl) that yields robust droplet formation (Fig. 1B). Indeed, for subsequent HXMS experiments (described, below), we sought to measure the behavior of an essentially homogenous droplet population since a highly heterogenous mixture of droplet and non-droplet ISB populations would likely yield convoluted mass spectra that would be challenging to properly assign to one of the states. In the condition we identified, 90% +/− 5% of the ISB protein was found within the rapidly sedimenting droplet population (Fig. 1C).

**Figure 1:**
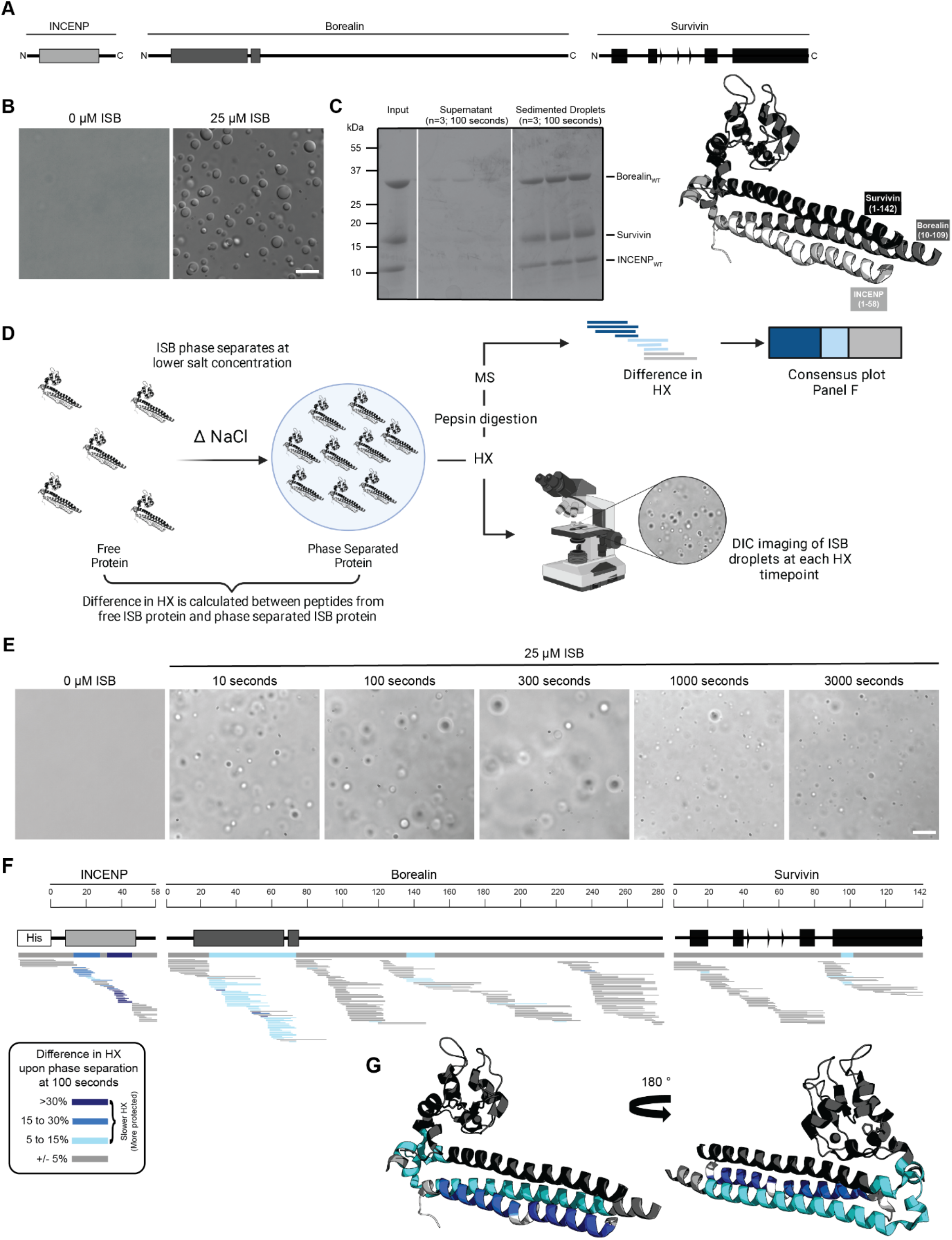
Phase separation leads to decreases in HX within the three-helix bundle structure of ISB. (A) Schematic of the ISB showing various structural domains within the subunits. Structural information was extracted from crystal structure of three-helix bundle structure of the ISB (PDB# 2QFA) ^16^. Each protein is color coded in a various shade of grey: INCENP = light grey, Borealin = mid-grey, Survivin = dark grey. (B) DIC micrographs of the ISB droplets under indicated conditions (25 µM ISB, 75 mM NaCl). Droplets were allowed to settle onto coverslip before imaging (∼5 minutes). Scale bar = 10 µm. (C) Sedimentation of pre-formed ISB droplets at phase separation conditions in Panel B (n=3). The time shown indicates the incubation period prior to sedimentation. (D) Schematic of HXMS experiments between free ISB protein and droplet ISB protein. HX samples either underwent pepsin digestion and analysis by MS or DIC imaging at each HX timepoint. Created with BioRender.com. (E) DIC micrographs of the ISB droplets at each HX timepoint (10s, 100s, 300s, 1000s, and 3000s). Droplets were not allowed to settle onto coverslip to allow for accurate timing of images. Scale bar = 10 µm. (F) Percent difference in HX is calculated for each peptide (represented by horizontal bars) at the 100 second time point and potted using the corresponding color key. The consensus behavior at each ISB residue is displayed in the horizontal bar below the secondary structure annotation taken from Panel A. These peptides were identified in a single experiment. When available, we present the data for all measurable charge states of the unique peptides within the experiment. (G) Consensus HXMS data from Panel F is mapped onto the three-helix bundle structure of the ISB, along with corresponding color key. Two views are shown, rotated by 180°.

We designed an HXMS experiment to compare the polypeptide backbone dynamics of the ISB in the free and droplet states (Fig. 1D). HXMS measures amide proton exchange, and for any generic protein, protection from HX is observed when secondary structures engage amide protons in hydrogen bonds ^17^. In our prior studies, we have utilized HXMS to readily identify contact points between domains of a multi-domain enzyme during its activation and inhibition ^18, 19^ as well as when components are added in a stepwise fashion during macromolecular complex assembly ^20, 21^. We reasoned that ISB backbone dynamics would be restricted upon droplet formation, since the generally accepted broadscale basis of phase separation is through intermolecular interactions, albeit transient ones. In the case of ISB, we assumed that inter-heterotrimer interactions were the basis of its subsequent phase separation. HXMS is routinely performed over a time course, and we developed an approach to monitor droplet formation behavior of the samples alongside the HX reactions themselves (Fig. 1D). Instead of letting the droplets settle on the slide, as in Fig. 1B, we monitored them as they exist immediately upon preparing the slides for imaging to provide a rapid readout of droplet formation at each timepoint (Fig. 1E). Robust droplet formation was observed in HX reaction conditions at all timepoints, including at the earliest one taken (10 s; Fig. 1E). By the latest timepoint, 3000 s, there was some diminution in the number of droplets (Fig. 1E), which may indicate the start of a transition of the droplets to a more solid state (i.e., gel-like). Thus, we concluded that timepoints longer than 3000 s would likely not be informative on how ISB backbone dynamics are impacted by initial droplet formation. This time course of HX proved to be sufficient to observe extensive exchange on all folded portions of the ISB, with the flexible regions lacking secondary structure exchanged much earlier (Supplementary Figs. 2 and 3). Slower HX was observed for all known and predicted secondary structural elements, except for the C-terminal helix of Borealin (Supplementary Fig. 2). Reciprocally, all predicted loop regions were very fast to exchange (i.e., essentially completely exchanged by 10 s), except for a small region around amino acids 140-150 of Borealin (Fig. 1F, Supplementary Figs. 4A-C). This region was originally interpreted to be largely unstructured and contain high amounts of intrinsic disorder; however, our HXMS analysis suggests that some secondary structural elements exist in this region and are central to phase separation. Notably, this region overlaps with a deletion mutant (Borealin^Δ139–160^) that causes a loss of phase separation ^8^.

To identify regions impacted by droplet formation, we first focused on an intermediate timepoint, 100 s, because visual inspection of HX patterns (Supplementary Fig. 2) indicated that there were clear changes at this point within the time course. At the 100 s timepoint, the most prominent differences between the soluble and droplet state were located within the three-helix bundle of the ISB, with long stretches in two subunits (INCENP and Borealin) and a small region at the N-terminal portion of the impacted α-helix in Survivin (Fig. 1F). The only other region that corresponded to slower HX was within the aforementioned region of Borealin (amino acids 140-150), displaying rates consistent with the presence of secondary structure when the ISB is in its free state, which became further accentuated within the droplet state (Fig. 1F). At the 300 s timepoint, a similar pattern is observed for the INCENP and Borealin proteins, with the notable addition of more extensive HX protection upon droplet formation within the three-helix bundle helix from Survivin and deprotection throughout its histone-recognizing BIR domain (Supplemental Figs. 4A, B, D). Taken together, the changes we observe in HX upon droplet formation indicates that discrete regions within structured portions of the ISB have slower backbone dynamics when in the droplet state.

### Two of Three Bundled ISB α-Helices Protected from HX in Droplets

We focused on the three prominent regions of the interacting α-helices of INCENP and Borealin. HX protection within INCENP is strongest at the 100 s time point, especially within the C-terminal portion of the α-helix (Figs. 2A-C). Examination of the entire time course shows that during intermediate levels of HX (i.e., between 100-1000 s), this region takes about three times as long to undergo the same amount of exchange when the ISB is in the droplet state relative to when it’s in the free protein state (Figs. 2B, C and Supplemental Fig. 2). Upon droplet formation, HX protection within Borealin is primarily located in the interacting α-helix and is less pronounced at any given peptide when compared to INCENP peptides (Fig. 2E). Nonetheless, similar to INCENP peptides, it still takes about twice as long to achieve the same level of deuteration for this region of Borealin in the droplet state as compared to the free state (Figs. 2F, G and Supplementary Fig. 2). In comparison, other regions exist within the ISB complex where multiple partially overlapping peptides show no measurable HX differences between droplet and free protein states (Fig. 2D). This verifies that there are no properties of droplets, such as vastly different molar concentrations of H_2_O (or D_2_O), that impact the general chemical exchange rate between all parts of the ISB. Rather, we conclude the changes observed within the ISB complex, such as those displayed in the two long interacting α-helices of INCENP and Borealin, are due to interactions formed between ISB complexes within the droplet state relative to those in the free state.

**Figure 2:**
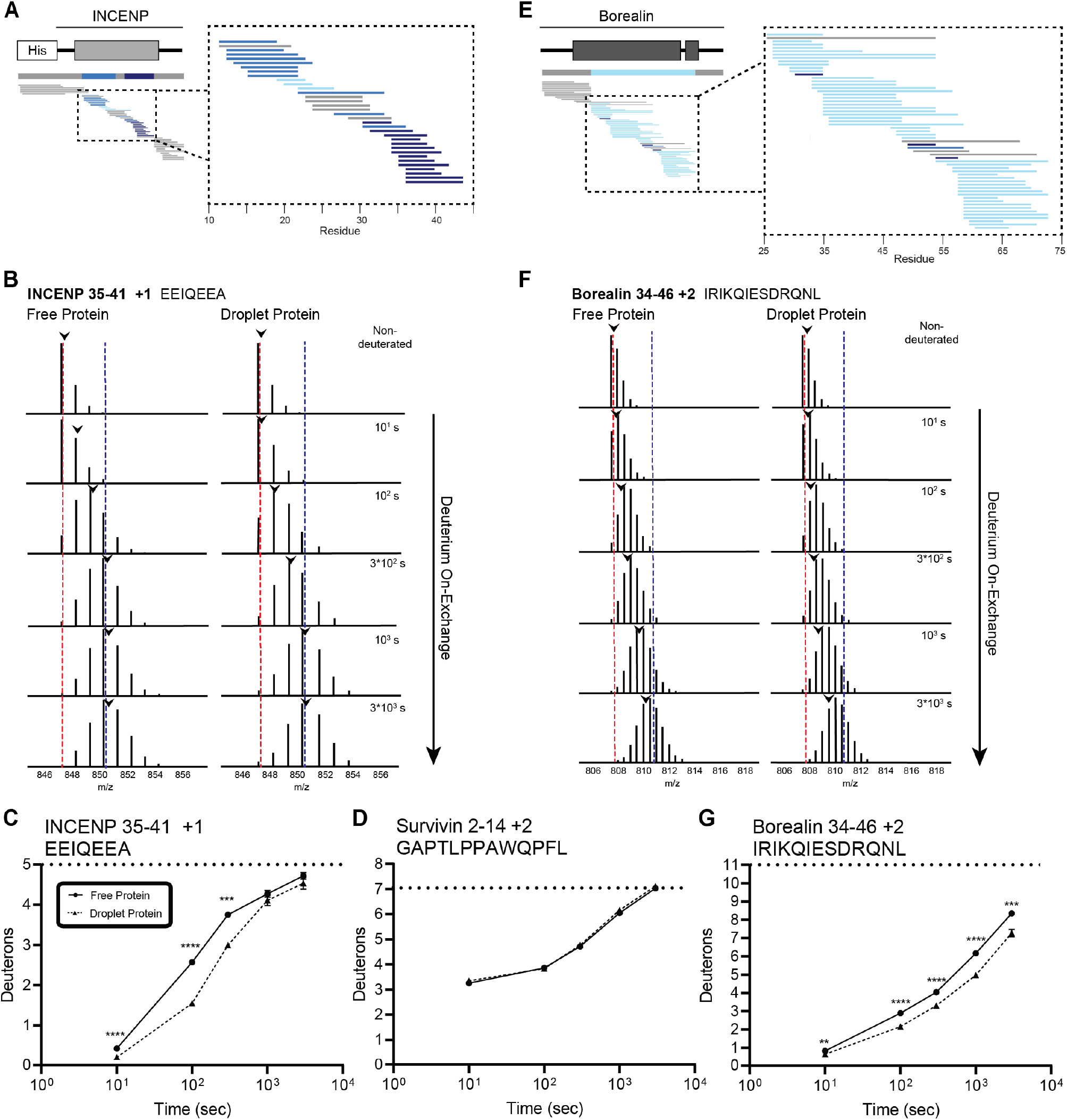
Regions of the three-helix bundle structure of INCENP and Borealin become protected from HX upon phase separation. (A) Percent difference in HX upon phase separation at 100 seconds in the indicated region of INCENP. (B) Raw MS data of a representative peptide from indicated region of INCENP. Centroid values are indicated with an arrowhead. Red and blue dotted lines serve as guides for visualizing differences. The red line lies on mono-isotopic peak whereas the blue line lies on the centroid value for the largest timepoint (3000 seconds) within the free protein sample. (C) HXMS of representative peptide from Panel B. The measured maximum number of exchangeable deuterons (maxD) when corrected with the average back exchange level (Supplementary Fig. 3B) is indicated. Data are represented as mean +/− s.e.m.; note: the error is too small to visualize outside of readable data points except in one instance. Statistical analysis was performed using multiple unpaired t-tests. **** p < 0.0001; *** 0.0001 < p < 0.001; ** 0.001 < p < 0.01. (D) HXMS of a peptide from the indicated region within Survivin and displayed, as described in panel C. This peptide shows the representative behavior of regions with the ISB that do not undergo changes in HX upon phase separation. Data are represented as mean +/− s.e.m.; note: the error is too small to visualize outside of readable data points. (E) Percent difference in HX upon phase separation at 100 seconds in the indicated region of Borealin. (F) Raw MS data of a representative peptide from indicated region of Borealin. Centroid values are indicated with an arrowhead. Red and blue dotted lines serve as guides for visualizing differences, as explained in Panel B. (G) HXMS of representative peptide from Panel F and displayed as described in panel C. Note: the error is too small to visualize outside of readable data points except in one instance.

### Phase Separation Involves an Acidic Surface Created by INCENP

We set to generate mutants to test the hypothesis that liquid-liquid demixing requires an interaction with the C-terminal portion of the long α-helix of INCENP within the three-helix bundle because this was the region with the greatest difference of HX protection between the droplet and free states (Fig. 2A). We anticipated an electrostatic component to ISB phase separation since droplet formation is restricted at higher ionic strength (Supplementary Fig. 1). A conspicuous stretch of glutamic acid residues (E35, E36, E39, E40, and E42; Fig. 3A) overlaps with a region of surface acidic charge in a single ISB heterotrimer ^16^. We found that mutation of all five glutamic acid residues to alanine (I_Mut1_SB; Fig. 3B) to remove the acidic charge within the region caused a visible reduction in large droplets relative to (ISB)_WT_ (Fig. 3C), although it was difficult to measure a difference using a standard turbidity (A_330_) measurement (Fig. 3D). Mutation of the five glutamic acid residues to arginine (I_Mut2_SB; Fig. 3B) to reverse the charge yielded a predictably more pronounced effect, observed in both the microscope-based and turbidity assessments (Figs. 3C, D). We conclude that some or all the five INCENP glutamic acid residues are involved in ISB phase separation.

**Figure 3:**
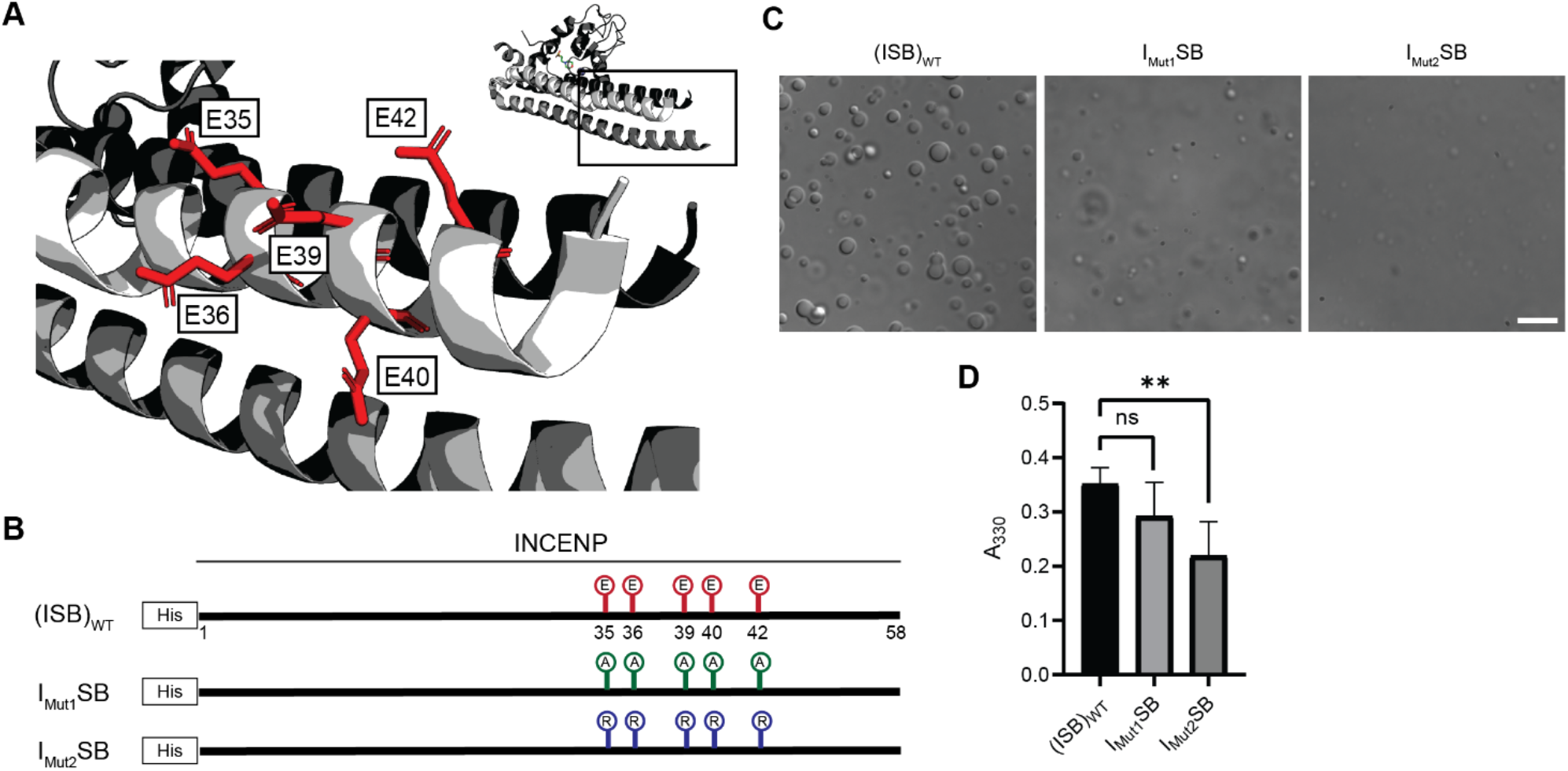
Acidic patch within INCENP coiled-coiled region contributes to electrostatic interaction within droplets. (A) Location of indicated acidic residues (E35/36/39/40/42) within INCENP at the surface of the coiled-coiled structure. Side chains are colored in red to indicate acidic charge. (B) Summary of a first round of mutations made to acidic residues within INCENP. Lolli-pop sticks represent each of the five residues in question. For (ISB)_WT_, red color indicates acidic charge. For I_Mut1_SB, green color indicates neutral charge. For I_Mut2_SB, blue color represents basic charge. (C) DIC micrographs of the ISB droplets for I_Mut1_SB and I_Mut2_SB. Scale bar = 10 µm. (D) Turbidity calculations of I_Mut1_SB and I_Mut2_SB measured as absorbance at 330 nm; n= 6 for (ISB)_WT_, I_Mut1_SB and I_Mut2_SB. Statistical analysis was performed using a Brown-Forsythe and Welch ANOVA test. ** 0.001 < p < 0.01.

### Crystal Packing-guided Mutagenesis to Disrupt Phase Separation

Liquid-liquid phase separation has been long studied as involving on- or off-pathway nucleation of a crystal lattice, occurring in different parts of the same phase diagram ^22^.Therefore, we assessed the crystal packing of ISB heterotrimers, and found that INCENP and Borealin from separate ISB heterotrimers are in close contact (PDB#2QFA) ^16^. Indeed, there were interactions between the HX protected regions of INCENP and Borealin (Figs. 2A, E), including several sidechains that we hypothesized to be involved in complementary electrostatic interactions (Fig. 4A). Specifically, we noted three acidic INCENP residues mutated in I_Mut2_SB were within salt bridge distance (i.e., ∼2-4 Å) from a corresponding positive residue on Borealin (Fig. 3). We used this as the basis for a second round of mutagenesis (Figs. 4B-D). Mutation of two of the acidic INCENP residues (E36 and E40; I_Mut3_SB) led to a similar level of reduction in phase separation (Figs. 4C, D) as when all five of the original glutamic acidic residues were mutated (Fig. 3). The addition of either one (E36, E40, and E42; I_Mut4_SB) or two (D27, E36, E40, and E42; I_Mut5_SB) mutated acidic INCENP residues displayed either similar or increased levels of reduction, respectively (Figs. 4C, D). These findings, along with our HX measurements (Fig. 1), raised the possibility of shared interaction sites between liquid-liquid demixed and crystal forms of the ISB.

**Figure 4:**
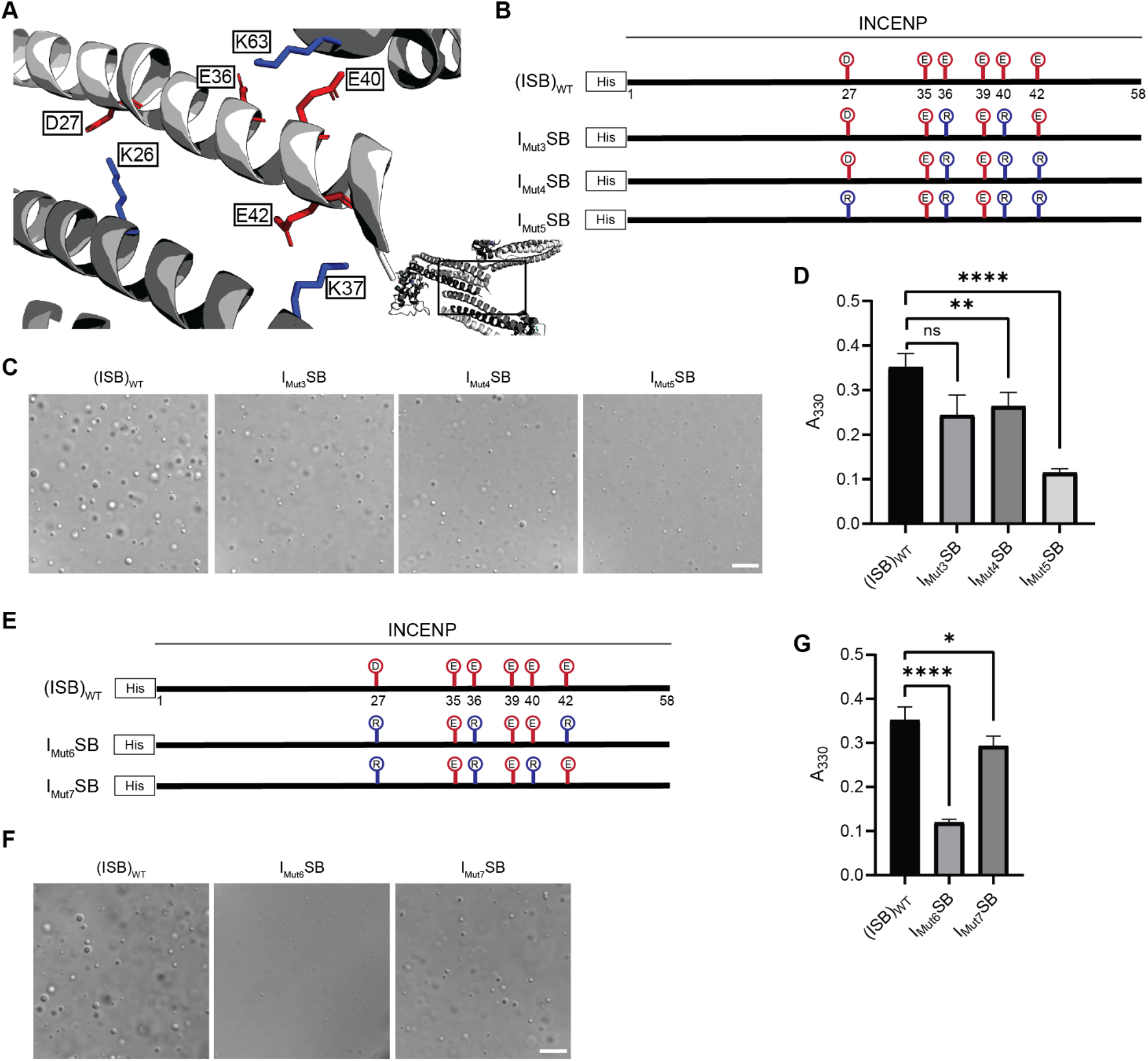
Crystal packing of ISB three-helix bundle structure highlights possible salt-bridges between multiple complexes. (A) Location of acidic and basic residues within crystal packing of ISB between INCENP_1_ and Borealin_2_ / Borealin_3_. Side chains are colored in red to indicate acidic charge and blue to indicate basic charge. (B) Summary of a second round of mutations made to acidic residues within INCENP. Lolli-pop sticks represent each of the indicated residues in question. (C) DIC micrographs of the ISB droplets for I_Mut3_SB, I_Mut4_SB, and I_Mut5_SB. Scale bar = 10 µm (D) Turbidity calculations of I_Mut3_SB, I_Mut4_SB, and I_Mut5_SB measured as absorbance at 330 nm; n= 6 for (ISB)_WT_, I_Mut4_SB, and I_Mut5_Sb. n = 3 for I_Mut3_SB. Statistical analysis was performed using a Brown-Forsythe and Welch ANOVA test. **** p < 0.0001; ** 0.001 < p < 0.01. (E) Summary of a third round of mutations made to acidic residues within INCENP. Lolli-pop sticks represent each of the indicated residues in question. (F) DIC micrographs of the ISB droplets for I_Mut6_SB and I_Mut7_SB. Scale bar = 10 µm (G) Turbidity calculations of I_Mut6_SB and I_Mut7_SB measured as absorbance at 330 nm; n= 6 for (ISB)_WT_. n= 3 for I_Mut6_SB and I_Mut7_SB. Statistical analysis was performed using a Brown-Forsythe and Welch ANOVA test. **** p < 0.0001; * 0.01 < p < 0.05.

As a first test of this notion, we designed a third round of mutagenesis to probe any potential interactions of the acidic INCENP residues facing two different ISB trimers in the crystal lattice: INCENP^D27,E42^ contact the Borealin subunit of one heterotrimer, while INCENP^E36,E40^ contact another (Fig. 4A). We predicted that the contacts responsible for phase separation closely correspond to the highlighted crystal contacts. If so, then those within typical bond distances (i.e., ∼3-4 Å) would have a larger impact than those with distances in the crystal lattice that are too large to generate a salt-bridge. Along with mutation of INCENP^D27,E36^, I_Mut6_SB and I_Mut7_SB vary by either including a mutation of INCENP^E42^ (within salt-bridge distance [2.9Å] of Borealin^K37^; I_Mut6_SB; Supplementary Fig. 5) or a mutation of INCENP^E40^ (at too large a distance to bond with Borealin^K63^; I_Mut7_SB; Supplementary Fig. 5) (Fig. 4E). We found that I_Mut6_SB, wherein all three mutations impact salt-bridges, profoundly reduces phase separation (Figs. 4F, G). On the other hand, I_Mut7_SB only has a minor effect on phase separation (Figs. 4F, G). With the prior finding with I_Mut3_SB, we deduce that INCENP^E36^, but not INCENP^E40^, contributes to phase separation. Combining the information from the three rounds of mutagenesis, we conclude that residues D27, E36, and E42 of INCENP contribute additively to the disruption of phase separation we observe within I_Mut6_SB. Together, these findings provided an early indication that precise salt-bridges between ISB heterotrimers are key to its phase separation.

### Breaking and Reforming Salt-bridges to Modulate ISB Phase Separation

We set to test our prediction that multiple salt-bridges between ISB heterotrimers drive its phase separation by breaking the salt-bridges from the opposite subunit (Borealin) and by reconstituting the salt bridges through pairing each charge switch mutation with each other. A mutation to the Borealin subunit that contains the three relevant alanine to glutamic acid substitutions was designed based on the structural model and combined with either wild type INCENP (ISB_Mut_) or the Mut_6_ version of INCENP (I_Mut6_SB_Mut_) (Figs. 5A, B). Consistent with our prediction, ISB_Mut_ was severely crippled in its ability to undergo phase separation (Figs. 5C-E). Strikingly, the compensatory mutations in INCENP that reconstitute the salt bridges between ISB hetrotetramers completely restore droplet formation in I_Mut6_SB_Mut_, detectable by microscopy and spectroscopy (Figs. 5C, D), and almost entirely restores wild type behavior measured by the ISB concentration required to saturate droplet formation (Fig. 5E and Supplementary Fig. 6). The potent rescue when the two surfaces are simultaneously mutated provides clear support for the conclusion that salt-bridging between these parts of the structured portions of ISB drive phase separation.

**Figure 5:**
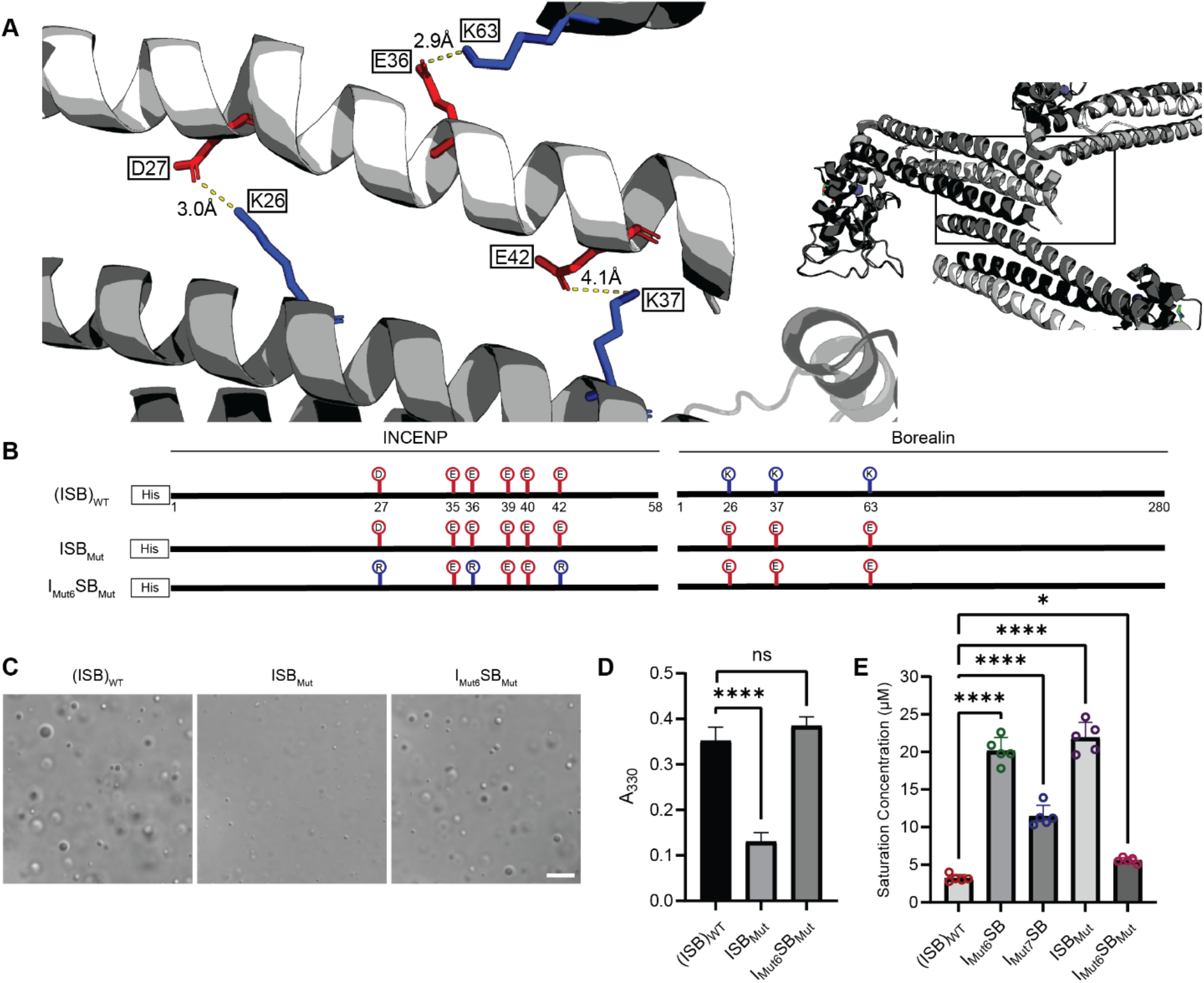
Salt-bridges between multiple ISB complexes provide multivalency required for phase separation. (A) Location of key salt-bridges within crystal packing of ISB between INCENP_1_ and Borealin_2_ / Borealin_3_. Side chains are colored in red to indicate acidic charge and blue to indicate basic charge. Distances between sidechains are indicated. (B) Summary of a fourth round of mutations made to acidic residues within INCENP and basic residues within Borealin. Lolli-pop sticks represent each of the indicated residues in question. (C) DIC micrographs of the ISB droplets for ISB_Mut_ and I_Mut6_SB_Mut._ Scale bar = 10 µm (D) Turbidity calculations of ISB_Mut_ and I_Mut6_SB_Mut_ at measured as absorbance at 330 nm; n= 6 for (ISB)_WT_. n= 3 for ISB_Mut_ and I_Mut6_SB_Mut_. Statistical analysis was performed using a Brown-Forsythe and Welch ANOVA test. **** p < 0.0001. (E) Saturation concentration of (ISB)_WT_, I_Mut6_SB, I_Mut7_SB, ISB_Mut_ and I_Mut6_SB_Mut_ in buffer containing 75 mM NaCl measured using sedimentation. n = 5 for (ISB)_WT_, I_Mut6_SB, I_Mut7_SB, ISB_Mut_ and I_Mut6_SB_Mut_. Statistical analysis was performed using a one-way ANOVA test with Dunnett’s multiple comparisons test. **** p < 0.0001; * 0.01 < p < 0.05.

### Mutation of Borealin to Disrupt Salt-bridges Reduces Phase Separation in Cells

To test whether or not the inter-CPC salt-bridges we identified impact phase separation in cells, we employed the Cry2 optoDroplet system ^23^, comparing Borealin_WT_ to Borealin_Mut_ (i.e. the same mutations present in ISB_Mut_ in Fig. 5). In this system, Borealin fused to Cry2, a light-inducible dimerizing protein, and mCherry (for fluorescent detection) readily forms droplets after exposing cells to blue light ^8^. We find that while Borealin_Mut_ can form droplets, their formation in the cytoplasm takes substantially longer than for Borealin_WT_ (Figs. 6A,B). Furthermore, the intensity of the mCherry signal in the droplets formed by Borealin_Mut_ is less than that observed with Borealin_WT_ (Figs. 6A,C). Together, these measurements indicate that mutating the salt-bridging residues we identified in Borealin (Figs. 4 and 5) reduces phase separation in the cellular environment (Fig. 6).

**Figure 6:**
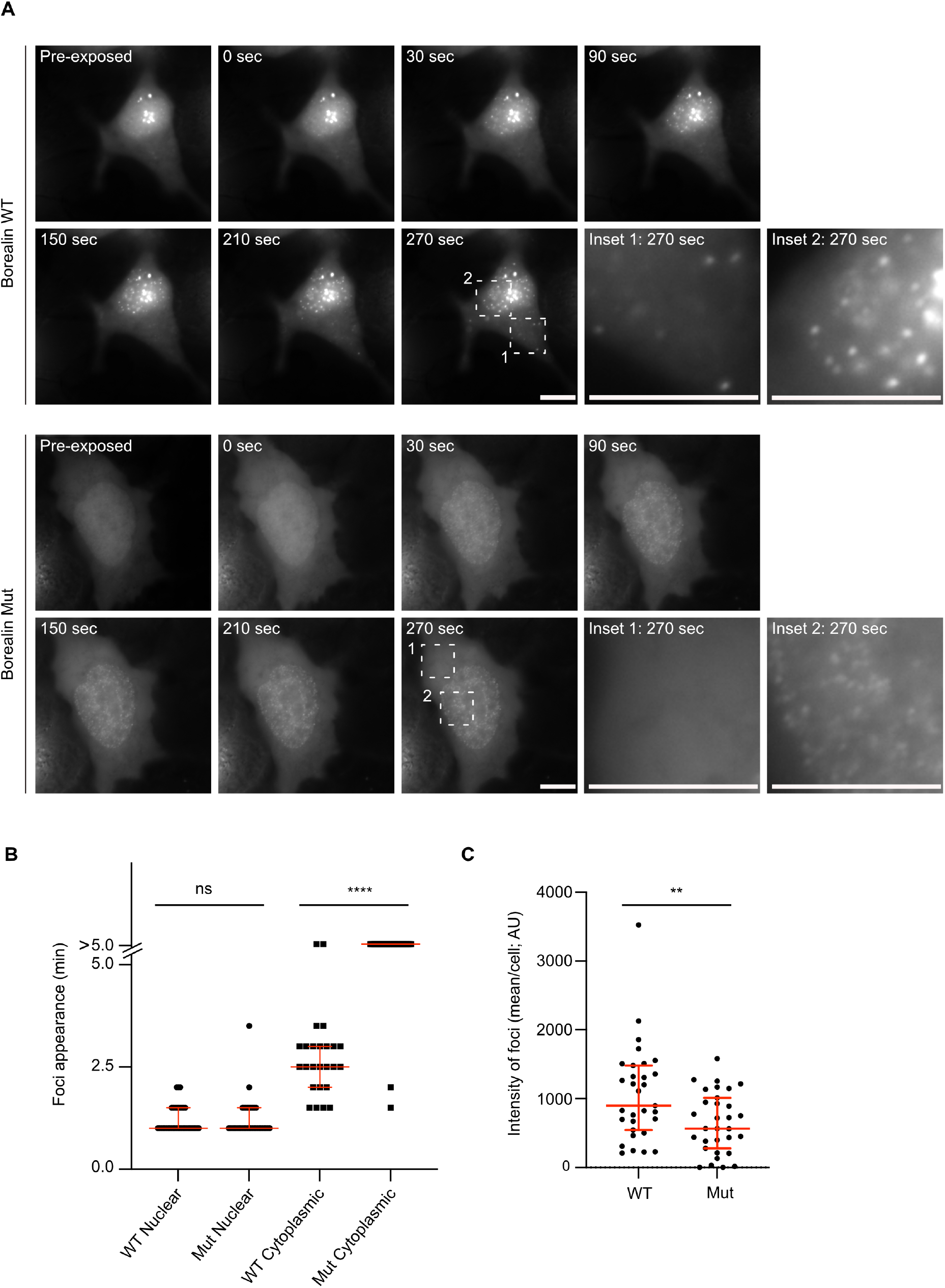
Disrupting salt-bridge residues in Borealin diminishes phase separation in cells. (A) Fluorescent detection of Borealin_WT_ or Borealin_Mut_, each fused to mCherry-Cry2 in an optoDroplet assay. Images were collected before and after (at the indicated timepoints) exposure to 488 nm light to induce Cry2 dimerization. Scale bar = 10 μm (B) Quantification of the time from induction of Cry2 dimerization until the appearance of foci. For each of the four types of measurements, 25-37 cells were measured, combined from two different experiments. The results of an unpaired, non-parametric t test, Mann-Whitney test are shown. **** p <0.0001. (C) Quantification of the intensity of foci. The results of an unpaired, non-parametric t test, Mann-Whitney test is shown, wherein ** equates to a p value, <0.01.

## Discussion

We demonstrate the feasibility of using HXMS as an unbiased approach to map the protein-protein interactions that drive liquid-liquid demixing. In the absence of structural information or useful structural models (i.e., of multimeric complexes where current structural predictions fall short), HXMS provides localization information at moderately high-resolution (i.e., within small numbers of amino acid residues given the coverage we achieved with the ISB, for instance). We envision that HXMS will be broadly useful to advance our physical understanding of the protein/protein and protein/nucleic acid interactions that drive phase separation and the formation of membraneless compartments within the cell.

It can be extremely difficult to generate structural information about the weak interactions that drive liquid-liquid demixing, but by combining HXMS with the long-studied relationship between liquid-liquid phase separation and crystal formation, our study shows that this can be done. Our approach might be generalizable for many proteins that have phase separation activities. Of course, packing information is largely thrown out during the presentation in most macromolecular structure-focused studies, but the data are readily available (i.e., archived in the PDB). We anticipate that clues from crystal contacts will help uncover potential sites that participate in other phase separating protein complexes, but they are unlikely to be sufficient. Rather, we envision HXMS as an essential step to localize the key contacts in the liquid-liquid demixed state, then most powerfully combined with structural information, including crystal contacts, when they are available.

Multivalency has emerged as the key driver of the liquid-liquid demixing of proteins and the interactions between monomeric units, and often involve weak interactions that are separated by intrinsically disordered domains. To date, detailed insight into the specific self-self interactions that are proposed to drive monomeric proteins and protein complexes into phase separated compartments has been rare, despite an enormous and growing list of candidate compartments at diverse locations in the cell ^1^. Undoubtedly, all sorts of macromolecular interaction types will be utilized within this diverse collection of compartments. Our findings support the notion that structured domains can play major roles as drivers of liquid-liquid demixing. Specifically, we map two structured regions that are separated by an intrinsically disordered region as underlying the liquid-liquid demixing of a subcomplex of the CPC. Our data suggest that salt-bridges between highly ordered regions of one subunit form with an adjacent complex in a liquid-liquid demixed droplet. These salt-bridges can explain the low affinity, salt sensitivity, and transient nature of the liquid demixed state. Our data also provide an important clue about the previously identified region on Borealin that is required for liquid-demixing in vitro and proper CPC assembly in cells ^8^. Specifically, our data (Fig. 1F, Supplementary Figs. 2, 4A) suggest this region of Borealin adopts secondary structure that undergoes additional HX protection in the liquid-liquid demixed state. We presume this region, enriched in basic residues, interacts with a negatively charged region that remains to be identified in an adjacent heterotrimer. We note that there is a region of strong negative charge within the intrinsically disordered region between the two regions of HX protection on the Borealin subunit. Our findings integrate into the emerging concept of ‘stickers and spacers’ ^24^, where the driving stickers for the CPC are indeed the specific salt-bridges that we identified through a combination of HXMS, analysis of ISB crystal packing, and mutagenesis. The spacers include the regions that lack stable secondary structure (i.e., the substantial portions where HX is essentially complete by the 10 s timepoint; Supplementary Fig. 2). The stickers are sufficient for higher order assembly, and the spacers permit particular solvation properties that contribute to the nature of the phase separation properties ^24–27^ and downstream consequences to the viscoelastic properties of the inner centromere compartment (Fig. 7).

**Figure 7:**
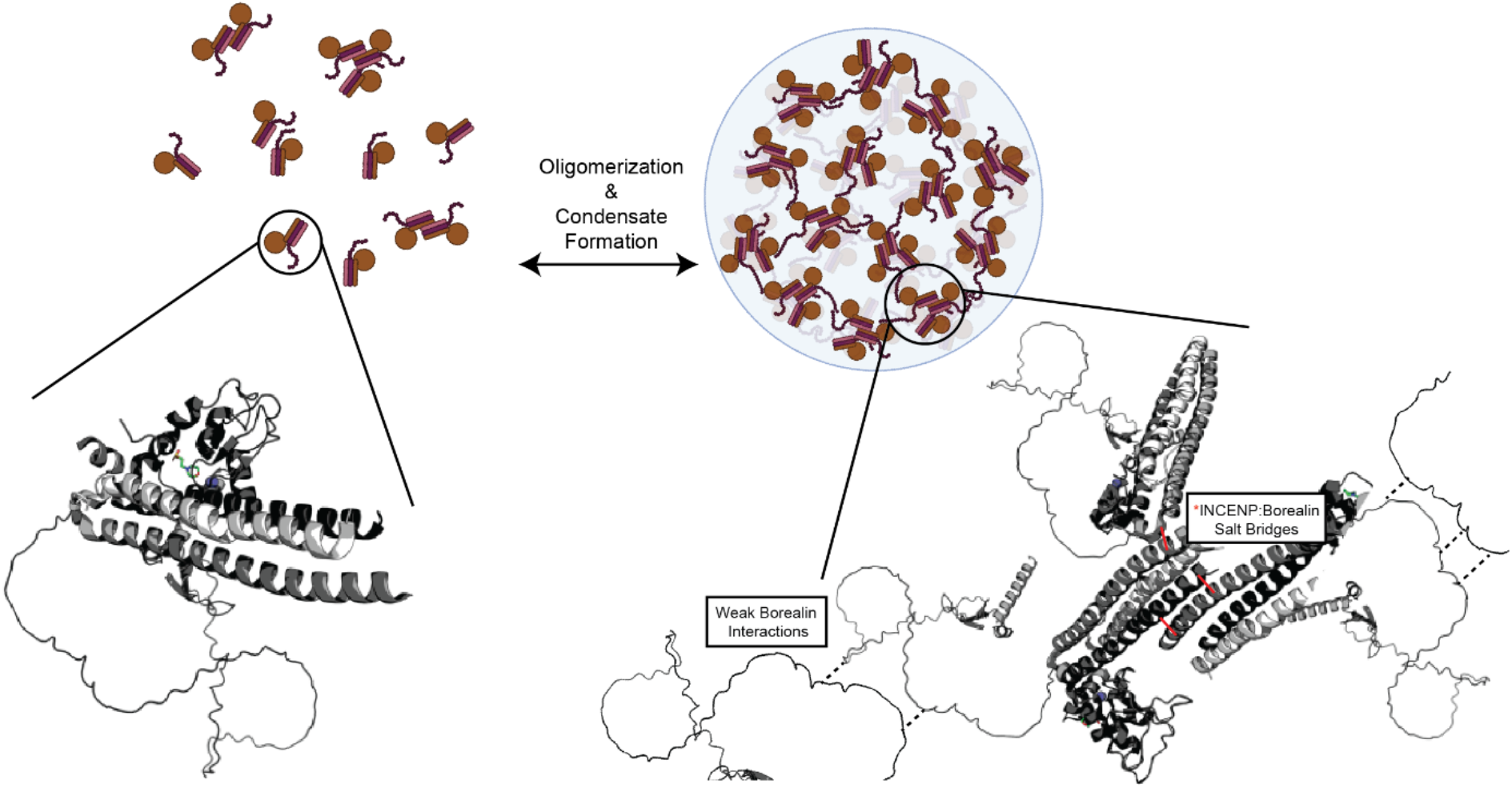
Summary model highlighting functionality of CPC phase separation in cells. Stabilized interactions defined by HXMS findings are indicated in solid black lines, while proposed weak interactions via Borealin loop are defined by dashed line. This model utilizes an AlphaFold prediction for the unstructured region of Borealin ^32, 33^. Created with BioRender.com.

Inner centromere formation is highly regulated so that the non-membranous compartment only forms during prophase and can be quickly disassembled at the beginning of anaphase ^28^. The reaction requires a combination of specific recruitment of the CPC to histone posttranslational modifications ^11–13^, interactions of the Borealin subunit with histones, and interactions between Borealin and the Sgo1 protein, which in turn is recruited by histone phosphorylation in addition to the multivalent interactions between the INCENP, Survivin, and Borealin subunits that are the subject of this work. Although we initially suggested that recruitment relies upon initial recognition of H3^T3phos^ directly by Survivin ^11^ — and indirectly by interaction with the Sgo2 adaptor protein that recognizes H2A^T120phos^ ^13^ — which then drives liquid demixing, it is also possible that the system is so finely tuned that the CPC can both interact with phosphohistones and also interact with other CPCs by multivalency, generating a chromosome-localized compartment capable of efficiently mediating robust mitotic error correction.

It has long been appreciated that Aurora B, the catalytic subunit of the CPC, is a prime example of a kinase whose activity is dictated by localization ^29^. We focus on the inner centromere, where Aurora B monitors connections of chromosomes to the mitotic spindle, and it is also vital later in mitosis for the abscission checkpoint and the regulation of cytokinesis ^30^. Since perturbing the ability of the CPC to phase separate at the inner centromere leads to defects in mitosis ^8, 31^, the role of phase separation appears to be tuning levels at the centromere, producing a functional inner centromere compartment, and maintaining a pool of the CPC after chromosome alignment destined for downstream steps in cytokinesis. In the present study, we have defined the impact of phase separation on the structural dynamics of the responsible regions for separation of the CPC, and, further, identified the key structural determinants that drive the inter-heterotrimer interactions. Recapitulating the environment of the inner centromere with purified components will constitute a substantial future challenge for the field, and our work provides a framework with which to understand this critical region of mitotic chromatin and a powerful experimental approach, HXMS, with which to probe the protein components of the inner centromere.

## Acknowledgements

This work was supported by NIH grants GM130302 (B.E.B.) and GM134591 (N.W.B.). We acknowledge support of N.W.B. by the UPenn Structural Biology and Molecular Biophysics Training Grant (GM008275). J. Shorter (UPenn) provided guidance on phase separation experiments.

## Author Contributions

N.W.B., P.T.S., and B.E.B. devised the project. N.W.B. performed all of the experiments except for the Cry2 optoDroplet experiments, which were performed by A.A. N.W.B. and E.N. analyzed crystallography data in light of HXMS results. L.M. assisted with HXMS experiments. N.W.B., L.M., and B.E.B. analyzed data. N.W.B. and B.E.B. wrote the paper. All authors edited the paper. B.E.B. directed the research.

## Data Availability

Source data are provided with this paper. The HXMS data in this study has been deposited in the Pride database under accession code PXD034374. The structure 2QFA [https://doi.org/10.2210/pdb2QFA/pdb] from the Protein Data Bank (www.rcsb.org) was used in this study. An AlphaFold prediction for the Borealin protein (primary accession number Q53HL2) was used in this study.

## Ethics Declarations

The authors declare no competing interests.

## Materials and Methods

### Protein Purification

Rosetta 2 (DE3) pLysS cells were transformed with a triscistronic pET28a vector containing sequences for 6xHis-INCENP^1–58^, FL survivin and FL boreain. Cells were then grown in the presence of 30 µg ml^−1^ kanamycin to an optical density (OD) between 0.6-0.8 and protein expression was induced with 1 mM isopropylthiogalactoside (IPTG) for 16-18 hours at 18⁰ C. The medium was also supplemented with 60 mg l^−1^ ZnCl_2_ and 0.2% glucose. Cells were then pelleted and lysed in buffer containing 50 mM Tris pH 7.5, 500 mM NaCl, 5% glycerol, 5 mM imidazole, 5 mM 2-mercaptoethanol (BME) and protease inhibitor cocktail (Roche) using a combination of dounce homogenization and sonication. The lysate was then cleared by centrifugation and purified over HisTrap HP column (Cytiva) and eluted using 50 mM Tris pH 7.5, 500 mM NaCl, 5% glycerol, 500 mM imidazole, 5 mM BME at 4⁰ C. The elutate was further gel filtered over a Hi-Load 16/60 Superdex-200 pg column (GE Life Sciences, Cytiva) in buffer containing 50 mM Tris pH 7.5, 500 mM NaCl, 5% glycerol and 5 mM BME. The desired fractions were collected and concentrated using Amicon Ultra-4 Centrifugal Filter Units with 3 kDa cut-off. All mutants within this study are purified similarly (Supplemental Fig. 7).

### Plasmid Construction and Mutagenesis

The I_Mut1_SB, I_Mut2_SB, I_Mut3_SB, I_Mut4_SB, and I_Mut5_SB constructs were created by a two-fragment assembly system (NEB), replacing WT INCENP^1–58^ sequence with the corresponding gBlock Gene Fragments (IDT). WT template DNA was amplified via PCR (Forward Primer: 3’ - TGAGATCCGAATTCGAGCTCTAATTTTG - 5’, Reverse Primer: 3’ - GCTGTGATGATGATGATGATGGCTGCTG) before assembly. The I_Mut6_SB construct was created by the Quikchange protocol (Stratagene) (Forward Primer: 3’ - CTTGAGCGTATCCAAGAGGAGGCCCGACGCATGTTCACC - 5’, Reverse Primer: 3’ - GGTGAACATGCGTCGGGCCTCCTCTTGGATACGCTCAAG - 5’), using the I_Mut5_SB construct as the template DNA. The I_Mut6_SB construct was created by Quikchange protocol (Stratagene) (Forward Primer: 3’ - CGTATCCAAGAGCGAGCCGAGCGCATGTTCACCAGAGAA - 5’, Reverse Primer: 3’ - TTCTCTGGTGAACATGCGCTCGGCTCGCTCTTGGATACG - 5’), using the I_Mut5_SB construct as the template DNA. The ISB_Mut_ and I_Mut6_SB_Mut_ constructs were created by a two-fragment assembly system (NEB), replacing WT INCENP^1–58^ and WT Borealin sequence with the corresponding gBlock Gene Fragment (IDT). WT template DNA was amplified via PCR (Forward Primer: 3’ - CCGTCTCGCCCAAATCTGCA - 5’, Reverse Primer: 3’ - GCTGTGATGATGATGATGATGGCTGCTG - 5’) before assembly. Sequences were verified by automated cycle sequencing (University of Pennsylvania Genomics Analysis Core).

I_Mut1_SB G Block: 3’ - CAG CAG CCA TCA TCA TCA TCA TCA CAG CAG CGG CCT GGT GCC GCG CGG CAG CCA TAT GGG GAC GAC GGC CCC AGG GCC CAT TCA CCT GCT GGA GCT ATG TGA CCA GAA GCT CAT GGA GTT TCT CTG CAA CAT GGA TAA TAA GGA CTT GGT GTG GCT TGC TGC CAT CCA AGC CGC AGC CGC TCG CAT GTT CAC CAG AGA ATT CAG CAA AGA GCC AGA GCT GAT GCC CAA ATG AGA TCC GAA TTC GAG CTC TAA TTT TG - 5’

I_Mut2_SB G Block: 3’ - CAG CAG CCA TCA TCA TCA TCA TCA CAG CAG CGG CCT GGT GCC GCG CGG CAG CCA TAT GGG GAC GAC GGC CCC AGG GCC CAT TCA CCT GCT GGA GCT ATG TGA CCA GAA GCT CAT GGA GTT TCT CTG CAA CAT GGA TAA TAA GGA CTT GGT GTG GCT TCG TCG AAT CCA ACG TCG AGC CCG TCG CAT GTT CAC CAG AGA ATT CAG CAA AGA GCC AGA GCT GAT GCC CAA ATG AGA TCC GAA TTC GAG CTC TAA TTT TG - 5’

I_Mut3_SB G Block: 3’ - CAG CAG CCA TCA TCA TCA TCA TCA CAG CAG CGG CCT GGT GCC GCG CGG CAG CCA TAT GGG GAC GAC GGC CCC AGG GCC CAT TCA CCT GCT GGA GCT ATG TGA CCA GAA GCT CAT GGA GTT TCT CTG CAA CAT GGA TAA TAA GGA CTT GGT GTG GCT TGA GCG TAT CCA AGA GCG AGC CGA GCG CAT GTT CAC CAG AGA ATT CAG CAA AGA GCC AGA GCT GAT GCC CAA ATG AGA TCC GAA TTC GAG CTC TAA TTT TG - 5’

I_Mut4_SB G Block: 3’ - CAG CAG CCA TCA TCA TCA TCA TCA CAG CAG CGG CCT GGT GCC GCG CGG CAG CCA TAT GGG GAC GAC GGC CCC AGG GCC CAT TCA CCT GCT GGA GCT ATG TGA CCA GAA GCT CAT GGA GTT TCT CTG CAA CAT GGA TAA TAA GGA CTT GGT GTG GCT TGA GCG TAT CCA AGA GCG AGC CCG ACG CAT GTT CAC CAG AGA ATT CAG CAA AGA GCC AGA GCT GAT GCC CAA ATG AGA TCC GAA TTC GAG CTC TAA TTT TG - 5’

I_Mut5_SB G Block: 3’ - CAG CAG CCA TCA TCA TCA TCA TCA CAG CAG CGG CCT GGT GCC GCG CGG CAG CCA TAT GGG GAC GAC GGC CCC AGG GCC CAT TCA CCT GCT GGA GCT ATG TGA CCA GAA GCT CAT GGA GTT TCT CTG CAA CAT GCG TAA TAA GGA CTT GGT GTG GCT TGA GCG TAT CCA AGA GCG AGC CCG ACG CAT GTT CAC CAG AGA ATT CAG CAA AGA GCC AGA GCT GAT GCC CAA ATG AGA TCC GAA TTC GAG CTC TAA TTT TG - 5’

ISB_Mut_ G Block: 3’ - CAG CAG CCA TCA TCA TCA TCA TCA CAG CAG CGG CCT GGT GCC GCG CGG CAG CCA TAT GGG GAC GAC GGC CCC AGG GCC CAT TCA CCT GCT GGA GCT ATG TGA CCA GAA GCT CAT GGA GTT TCT CTG CAA CAT GGA TAA TAA GGA CTT GGT GTG GCT TGA GGA AAT CCA AGA GGA GGC CGA GCG CAT GTT CAC CAG AGA ATT CAG CAA AGA GCC AGA GCT GAT GCC CAA ATG AGA TCC GAA TTC GAG CTC TAA TTT TGT TTA ACT TTA AGA AGG AGA TAT ACC ATG GCT CCT AGG AAG GGC AGT AGT CGG GTG GCC AAG ACC AAC TCC TTA CGG AGG CGG AAG CTC GCC TCC TTT CTG GAG GAC TTC GAC CGT GAA GTG GAA ATA CGA ATC GAG CAA ATT GAG TCA GAC AGG CAG AAC CTC CTC AAG GAG GTG GAT AAC CTC TAC AAC ATC GAG ATC CTG CGG CTC CCC GAG GCT CTG CGC GAG ATG AAC TGG CTT GAC TAC TTC GCC CTT GGA GGA AAC AAA CAG GCC CTG GAA GAG GCG GCA ACA GCT GAC CTG GAT ATC ACC GAA ATA AAC AAA CTA ACA GCA GAA GCT ATT CAG ACA CCC CTG AAA TCT GCC AAA ACA CGA AAG GTA ATA CAG GTA GAT GAA ATG ATA GTG GAA GAG GAA GAA GAA GAA GAA AAT GAA CGT AAG AAT CTT CAA ACT GCA AGA GTC AAA AGG TGT CCT CCA TCC AAG AAG AGA ACT CAG TCC ATG CAA GGA AAA GGA AAA GGG AAA AGG TCA AGC CGT GCT AAC ACT GTT ACC CCA GCC GTG GGC CGA TTG GAG GTG TCC ATG GTC AAA CCA ACT CCA GGC CTG ACA CCC AGG TTT GAC TCA AGG GTC TTC AAG ACC CCT GGC CTG CGT ACT CCA GCA GCA GGA GAG CGG ATT TAC AAC ATC TCA GGG AAT GGC AGC CCT CTT GCT GAC AGC AAA GAG ATC TTC CTC ACT GTG CCA GTG GGC GGC GGA GAG AGC CTG CGA TTA TTG GCC AGT GAC TTG CAG AGG CAC AGT ATT GCC CAG CTG GAT CCA GAG GCC TTG GGA AAC ATT AAG AAG CTC TCC AAC CGT CTC GCC CAA ATC TGC A - 5’

I_Mut6_SB_Mut_ G Block: 3’ - CAG CAG CCA TCA TCA TCA TCA TCA CAG CAG CGG CCT GGT GCC GCG CGG CAG CCA TAT GGG GAC GAC GGC CCC AGG GCC CAT TCA CCT GCT GGA GCT ATG TGA CCA GAA GCT CAT GGA GTT TCT CTG CAA CAT GCG TAA TAA GGA CTT GGT GTG GCT TGA GCG TAT CCA AGA GGA GGC CCG ACG CAT GTT CAC CAG AGA ATT CAG CAA AGA GCC AGA GCT GAT GCC CAA ATG AGA TCC GAA TTC GAG CTC TAA TTT TGT TTA ACT TTA AGA AGG AGA TAT ACC ATG GCT CCT AGG AAG GGC AGT AGT CGG GTG GCC AAG ACC AAC TCC TTA CGG AGG CGG AAG CTC GCC TCC TTT CTG GAG GAC TTC GAC CGT GAA GTG GAA ATA CGA ATC GAG CAA ATT GAG TCA GAC AGG CAG AAC CTC CTC AAG GAG GTG GAT AAC CTC TAC AAC ATC GAG ATC CTG CGG CTC CCC GAG GCT CTG CGC GAG ATG AAC TGG CTT GAC TAC TTC GCC CTT GGA GGA AAC AAA CAG GCC CTG GAA GAG GCG GCA ACA GCT GAC CTG GAT ATC ACC GAA ATA AAC AAA CTA ACA GCA GAA GCT ATT CAG ACA CCC CTG AAA TCT GCC AAA ACA CGA AAG GTA ATA CAG GTA GAT GAA ATG ATA GTG GAA GAG GAA GAA GAA GAA GAA AAT GAA CGT AAG AAT CTT CAA ACT GCA AGA GTC AAA AGG TGT CCT CCA TCC AAG AAG AGA ACT CAG TCC ATG CAA GGA AAA GGA AAA GGG AAA AGG TCA AGC CGT GCT AAC ACT GTT ACC CCA GCC GTG GGC CGA TTG GAG GTG TCC ATG GTC AAA CCA ACT CCA GGC CTG ACA CCC AGG TTT GAC TCA AGG GTC TTC AAG ACC CCT GGC CTG CGT ACT CCA GCA GCA GGA GAG CGG ATT TAC AAC ATC TCA GGG AAT GGC AGC CCT CTT GCT GAC AGC AAA GAG ATC TTC CTC ACT GTG CCA GTG GGC GGC GGA GAG AGC CTG CGA TTA TTG GCC AGT GAC TTG CAG AGG CAC AGT ATT GCC CAG CTG GAT CCA GAG GCC TTG GGA AAC ATT AAG AAG CTC TCC AAC CGT CTC GCC CAA ATC TGC A - 5’

### Phase-separation Assay

Phase separation was induced by diluting the indicated amount of ISB in a low salt buffer (50 mM Tris pH 7.5 and 5 mM BME) to achieve the indicated final concentration of protein and NaCl (25 µM ISB, 75 mM NaCl). Protein was always added last to each reaction. Phase separation was observed by adding a small volume of the reaction onto a coverslip and then imaging the ISB droplet by differential interference contrast (DIC) microscopy. All movies and images were captured within five minutes of the reaction set-up. For time-lapse imaging of ISB droplet fusion, ISB droplets were formed in the indicated conditions and immediately imaged via DIC every second. Imaging of the ISB droplet during HXMS experimentation was captured as close to the indicated time-point as possible.

### Sedimentation Assay

Following liquid-liquid phase separation, each reaction was allowed to stand for 100 seconds and then centrifuged at 16,100 ᵡ g for 10 minutes to separate the soluble phase from the droplet phase. All of the top phase was removed and placed in a separate tube. The dense phase was resuspended in an equivalent volume of purification buffer (50 mM Tris pH 7.5, 500 mM NaCl, 5% glycerol, 5 mM BME). Then, 10 µL from each top phase and dense phase was removed and analyzed using SDS-PAGE.

### HXMS measurement and analysis

Deuterium on-exchange of soluble ISB protein was performed at room temperature (25°C) by diluting purified ISB with deuterium on-exchange buffer (50 mM Tris pD 7.5, 500 mM NaCl) to a final protein concentration of 25 µM ISB, 500 mM NaCl and a final D_2_O content of 75%. A 20 µL aliquot was removed at each time point (10, 100, 300, 1000, 3000 seconds) and the reaction was quenched with 30 µL ice-cold quench buffer (1.67 M guanidine hydrochloride, 8% glycerol, and 0.8% formic acid, for a final pH of 2.4-2.6) and rapidly frozen in liquid nitrogen. The samples were stored at −80⁰ C until analysis by MS. Deuterium on-exchange of phase-separated ISB protein was performed at a similar temperature by diluting purified ISB with a mixture of two on-exchange buffers (Buffer 1: 50 mM Tris pD 7.5, 0 mM NaCl; Buffer 2: 50 mM Tris pD 7.5, 500 mM NaCl) to a final protein concentration of 25 µM ISB, 75 mM NaCl and a final D_2_O content of 75%. pD values are direct pH meter readings. Samples were prepared and frozen in a similar manner to soluble ISB protein. All samples were produced in quadruplicate so that there would be a spare in addition to a triplicate set to measure, in case of a technical issue in downstream steps. The supplementary table summarizes the HXMS experiments HX samples were individually thawed at 0⁰ C for 2.5 minutes, then injected (50 µL) and pumped through an immobilized pepsin (Sigma) column at an initial flow rate of 50 µL/min for 2 minutes followed by 150 µL/min for 2 minutes. Pepsin was immobilized by coupling to POROS 20 AL support (Applied Biosystems) and packed into column housings of 2mm x 2cm (64 µL) (Upchurch). Protease-generated fragments were collected onto a TARGA C8 5 µM Piccolo HPLC column (1.0 x 5.0 mm, Higgins Analytical) and eluted through an analytical C18 HPLC column (0.3 x 75 mm, Agilent) by a shaped 12-100% buffer B gradient over 25 minutes at 6 µL/min (Buffer A: 0.1% formic acid; Buffer B: 0.1% formic acid, 99.9% acetonitrile). The effluent was electrosprayed into the mass spectrometer (LTQ Orbitrap XL, Thermo Fisher Scientific). We analyzed MS/MS data collected from ND samples to identify the likely sequence of the patent peptides using SEQUEST (Bioworks v3.3.1, Thermo Fisher Scientific) with a peptide tolerance of 8 ppm and a fragment tolerance of 0.1 AMU. We discarded peptides that failed to meet a specified quality score (SEQUEST P_pep_ score <0.99). The peptide data are included in the supplementary table.

A MATLAB based program, ExMS2 was used to prepare the pool of peptides based on SEQUEST output files. HDExaminer software was next used to process and analyze the HXMS data. HDExaminer identifies the peptide envelope centroid values for non-deuterated as well as deuterated peptides and uses the information to calculate the level of peptide deuteration for each peptide at each timepoint. Each individual deuterated peptide is corrected for loss of deuterium label during HXMS data collection by normalizing to the maximal deuteration level of that peptide, which we measure in a “full deuterated” (FD) reference sample. The FD sample was prepared in 75% deuterium to mimic the exchange experiment, but under acidic denaturing conditions (0.88% formic acid), and incubated for over 24 hours to allow each amide proton position along the entire polypeptide to undergo full exchange. 20 µL of this reaction was quenched with 30 µL ice-cold FD quench buffer (1M guanidine hydrochloride, 8% glycerol, and 0.74% formic acid, for a final pH of 2.4-2.6) and rapidly frozen in liquid nitrogen. HDExaminer performs such correction automatically when provided with the FD file. For each peptide, we compare the extent of deuteration as measured in both the on exchange and FD samples to the maximal number of exchangeable deuterons (maxD) when corrected with an average back exchange level; the median extent of back-exchange in our datasets is 18% (Supplementary Fig. 3).

### HXMS Plotting

Peptide plotting was performed in MATLAB, RStudio and Prism using deuteration levels for each peptide extracted from the HDExaminer outputs. Differences in deuteration levels between two samples were calculated for all peptides for which the identical peptide was found in both conditions, the ND and FD samples. For comparing two different HXMS datasets, we plot the percent difference of each peptide, which is calculated by subtracting the percent deuteration of one sample from that or another, and plotted according to the color legend in stepwise increments (as in Fig. 1F and Supplementary Fig. 4A). We include in our figures peptides of identical sequence but different charge states. Although not unique peptides, they do add confidence to our peptide identification as their deuteration levels are in close agreement with each other. Consensus behavior at each residue was calculated as the average of the differences in HX protection of all peptides spanning that residue (as in Fig. 1F and Supplemental Fig. 4A). For the plot of peptide data expressed as the number of deuterons (as in Figs. 2C-D and G, and Supplementary Figs. 4C-E), the values are expressed as the mean of three independent measurements +/− s.d.

### Turbidity Assay

Following liquid-liquid phase separation, WT and ISB mutant protein were incubated at room temperature for 100 seconds prior to UV-visible measurements. Control measurements included protein purification buffer, low salt buffer (50 mM Tris, 75 mM NaCl, 5% glycerol), and WT ISB protein at high salt (25 µM WT ISB, 500 mM NaCl). The optical intensity (turbidity) was measured using a Thermo Scientific NanoDrop 2000 UV-Vis spectrophotometer at 330 nm. The number of replicates is indicated in figure legends.

### Measuring Saturation Concentration

Following liquid-liquid phase separation, indicated proteins were incubated at room temperature for 100 seconds and centrifuged at 16,100 ᵡ g for 10 minutes to separate the soluble phase from the droplet phase. Then, the entirety of the top phase was removed; the remaining sedimented pellet was resuspended in an equivalent volume of protein purification buffer. 5 µL of both top phase and sedimented pellet, along with a sample of protein after thawing and a sample of protein before sedimentation, were analyzed using SDS-PAGE (4-20 % Tris-HCl gradient gel) to determine the saturation concentration. The serial dilution of wild-type ISB, ranging between 0 µM to 30 µM, was loaded onto a similar SDS-PAGE gel to create a standard curve (with a coefficient of determination R^2^=0.9), which was used to determine the saturation concentration of ISB. The SDS-PAGE gel was stained with Coomassie Blue and subjected to densitometry using GelQuantNET.

### OptoDroplet Assay

The plasmid expressing Borealin_WT_-mCherry-Cry2 ^8^ and a derivative harboring the K→E substitutions at Borealin a.a. 26, 37, and 63 were transfected in to HeLa TREx cells that were seeded in 35 mm glass bottom dishes (Cellvis, D29-20-1.5P). Lipofectamine 3000 (L3000-008) was used for transfections. Twenty-four hours following transfection, the cells were imaged using a Zeiss Observer-Z1 microscope in the presence of 5% CO_2_ in a humidified chamber at 37°C. Cells with similar mCherry expression levels were selected for measurement. To induce phase separation, the cells in the field were exposed to 488 nm light (10 cycles of 100 ms each, with an interval of 30 s between consecutive cycles). The single z-plane mCherry images were acquired immediately after exposure with 488 nm light for 1500 ms during each cycle. The cells were quantified for the time of appearance of mCherry foci in the nucleus and the cytoplasm. The mCherry intensities were quantified in Fiji (ImageJ) software and plotted using the GraphPad Prism software.

---

**Supplemental Figure 1:**
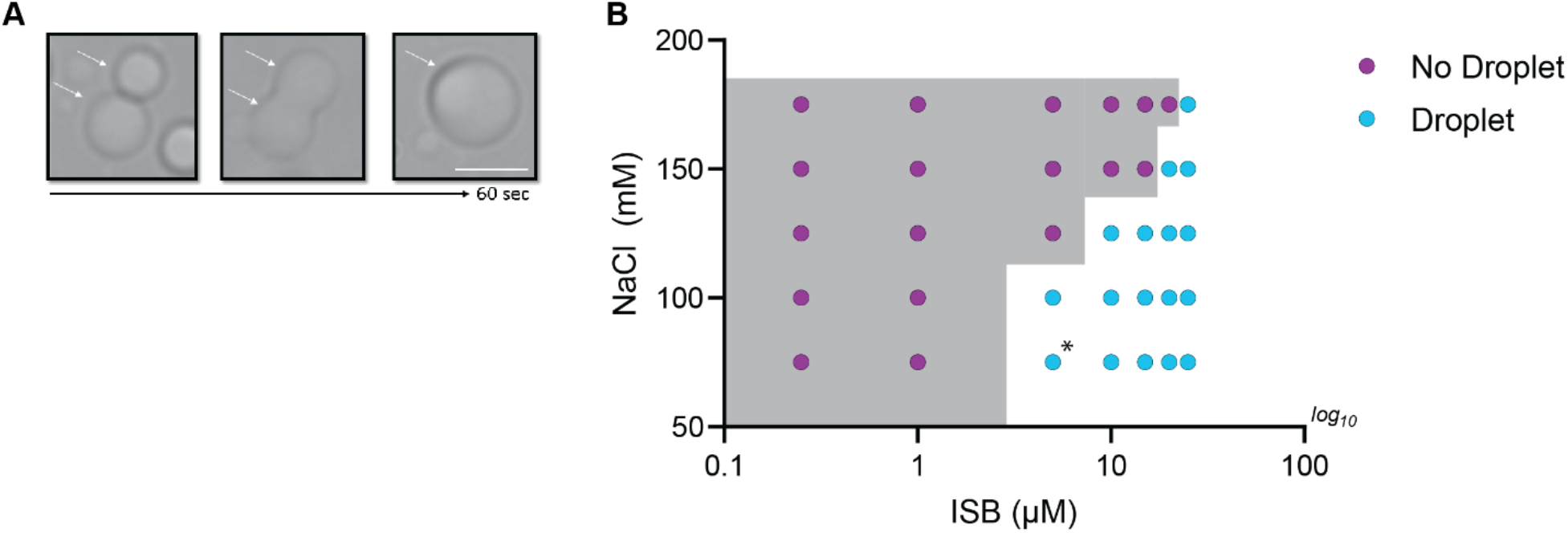
Phase properties of WT-ISB in absence of crowder. (A) Fusion of WT-ISB droplets as visualized by time-lapse imaging in the absence of crowding agent. (B) Phase diagram of WT-ISB phase separation as a function of the concentration of NaCl and ISB in the absence of crowding agent. The presence (cyan) or absence (magenta) of WT-ISB droplets. The grey area indicates the phase boundary. Conditions on either side of the phase boundary were repeated three times. For the condition [ISB] = 5 µM and [NaCl] = 75 mM, circle is highlighted with asterisk because two replicates displayed droplet formation, but one condition did not.

**Supplemental Figure 2:**
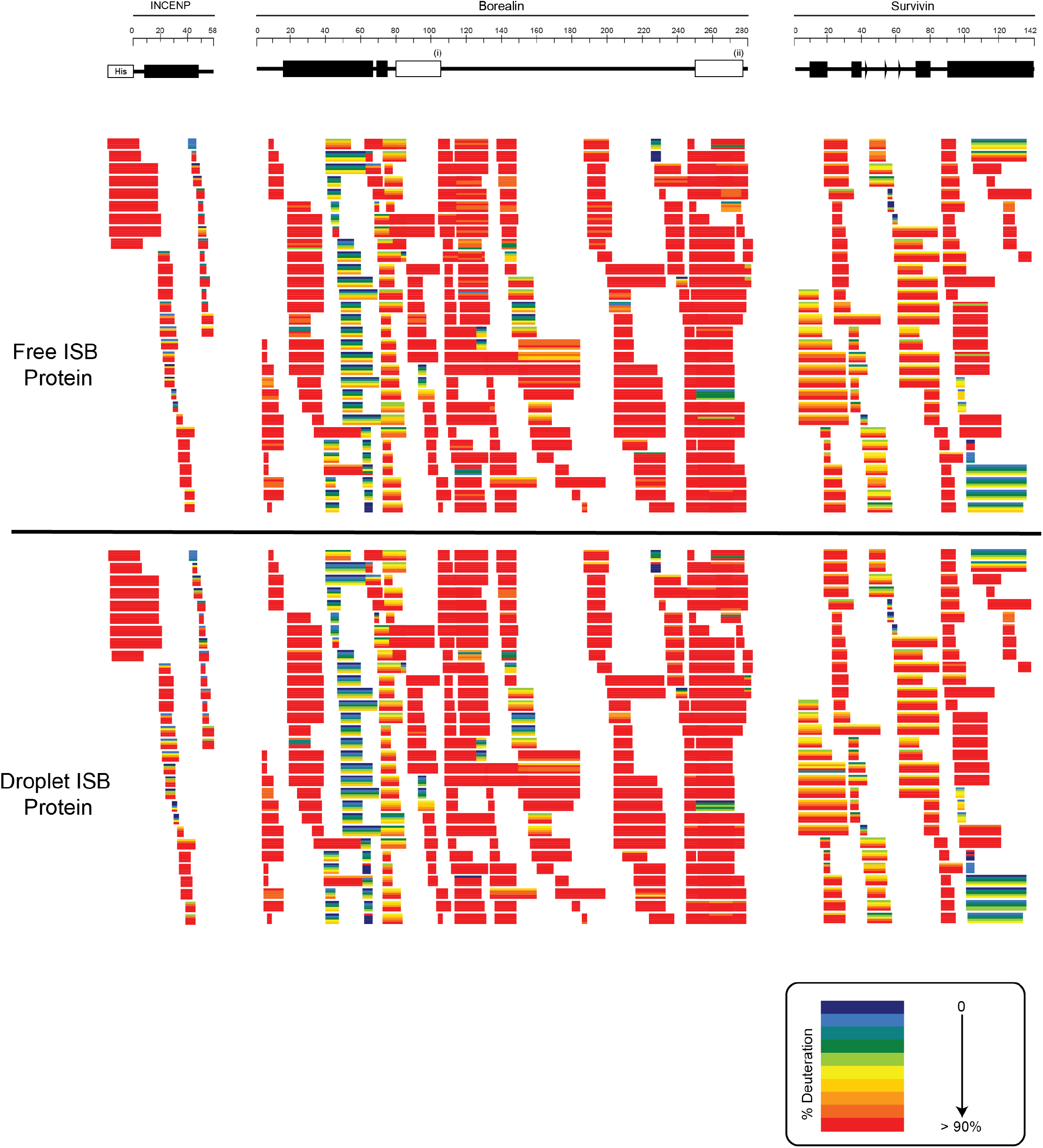
Ribbon Plots for Free and Droplet ISB Protein. HXMS data for free ISB Protein and droplet ISB Protein. Each horizontal bar represents an individual peptide, and the 5 stripes within each bar are colored according to the percentage deuteration at each of the 5 time points (10, 100, 300, 1000, 3000 s).

**Supplemental Figure 3:**
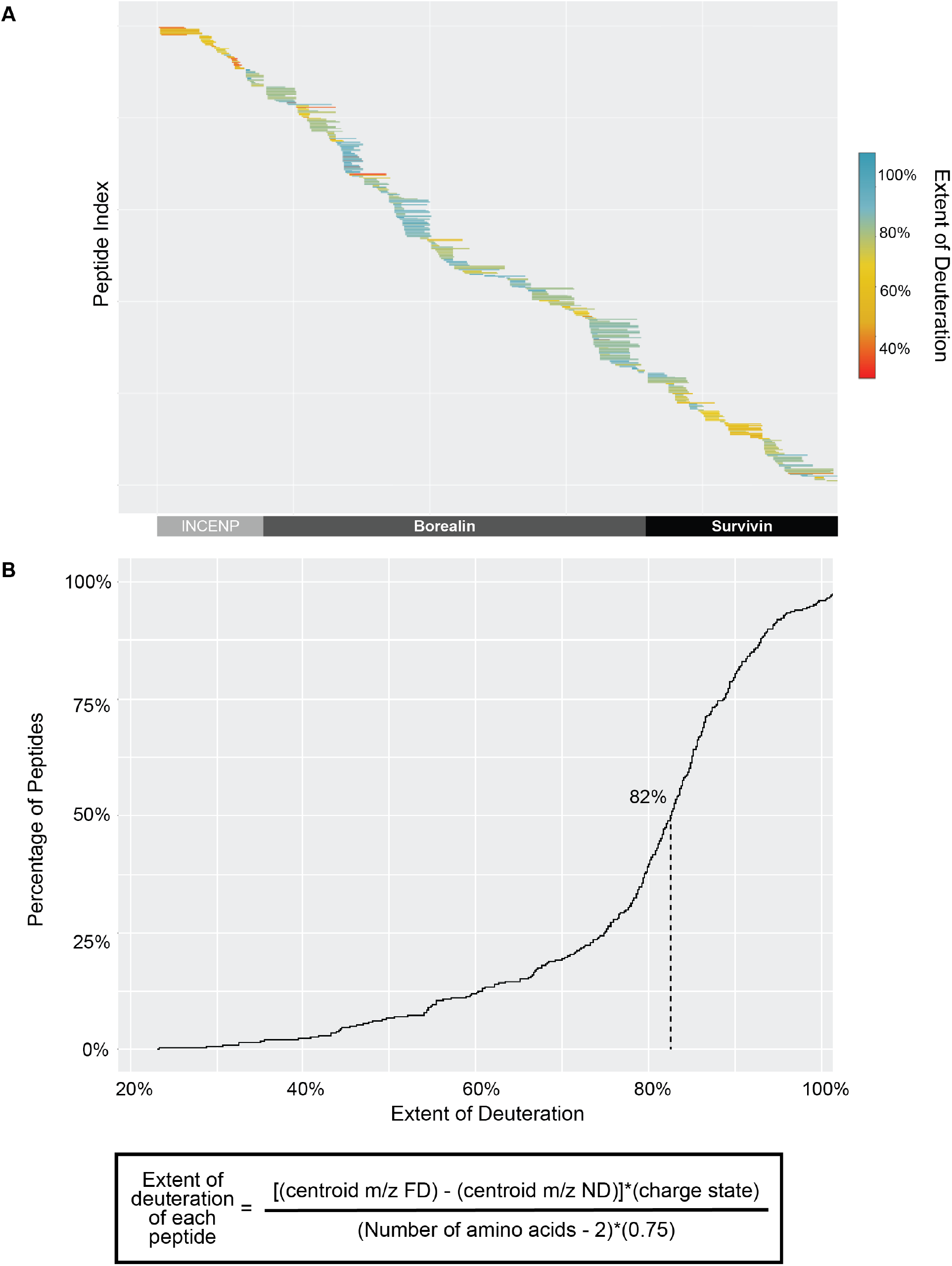
Extent of Deuteration within Fully Deuterated HXMS Control Samples. (A) Extent of peptide deuteration across INCENP, Borealin and Survivin sequence within a representative fully deuterated (FD) HXMS control samples. (B) Cumulative distribution curve of a representative fully deuterated (FD) sample, showing the extent of deuteration of all peptides compared to the theoretical maximum amount of deuteration of each peptide. The median deuteration was ∼82% for the FD sample, therefore the back-exchange after quench step was only ∼18%, which I well within optimal range.

**Supplemental Figure 4:**
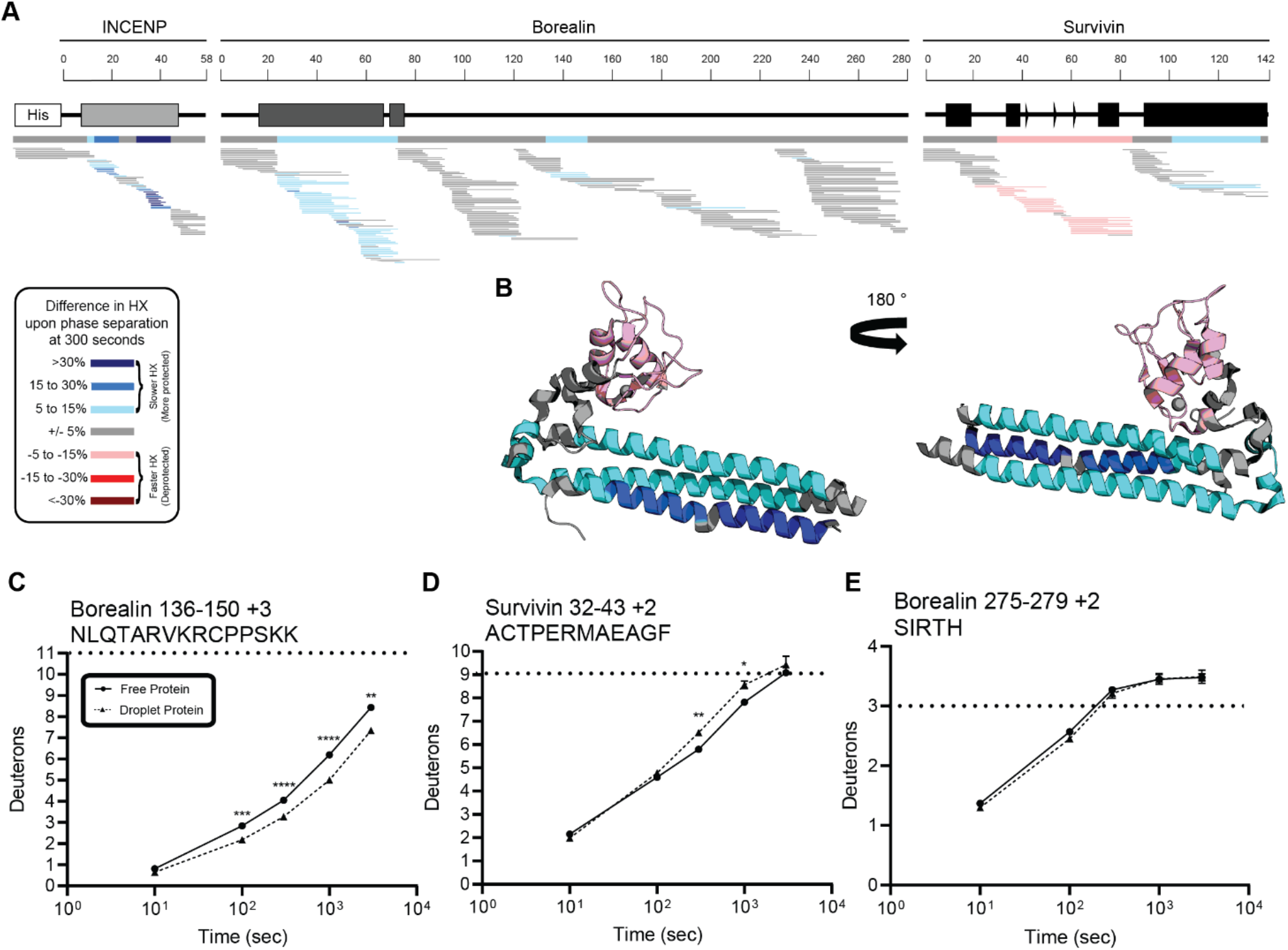
Percent difference in HX calculated for each peptide at 300 seconds. (A) Percent difference in HX is calculated for each peptide (represented by horizontal bars) at the 300 second time point and plotted using the corresponding color key. The consensus behavior at each ISB residue is displayed in the horizontal bar below the secondary structure annotation taken from Figure 1, Panel A. These peptides were identified in a single experiment. When available, we present the data for all measurable charge states of the unique peptides within the experiment. (B) Consensus HXMS data from Supplemental Figure 4A is mapped onto the three-helix bundle structure of the ISB. Two views are shown, rotated by 180°. (C) HXMS of representative peptide from indicated region of Borealin. Data are represented as mean +/− s.e.m.; note: the error is too small to visualize outside of readable data points. Statistical analysis was performed using multiple un-paired t-tests. **** p < 0.0001; *** 0.0001 < p < 0.001; ** 0.001 < p < 0.01. (D) HXMS of representative peptide from indicated region of Survivin and displayed as described in panel C. Data are represented as mean +/− s.e.m.; note: the error is too small to visualize outside of readable data points except in two instances. Statistical analysis was performed using multiple un-paired t-tests. ** 0.001 < p < 0.01; * 0.01 < p < 0.05. (E) HXMS of representative peptide from a region within Borealin and displayed as described in panel C. This peptide shows the representative behavior of regions with the ISB that do not undergo changes in HX upon phase separation. Data are represented as mean +/− s.e.m.; note: the error is too small to visualize outside of readable data points except in one instance. The apparent overcorrection, deuteration above the maxD level, is likely due to retained deuterium at position 2 due to the slowing effect of the Ilu side chain ^34^.

**Supplemental Figure 5:**
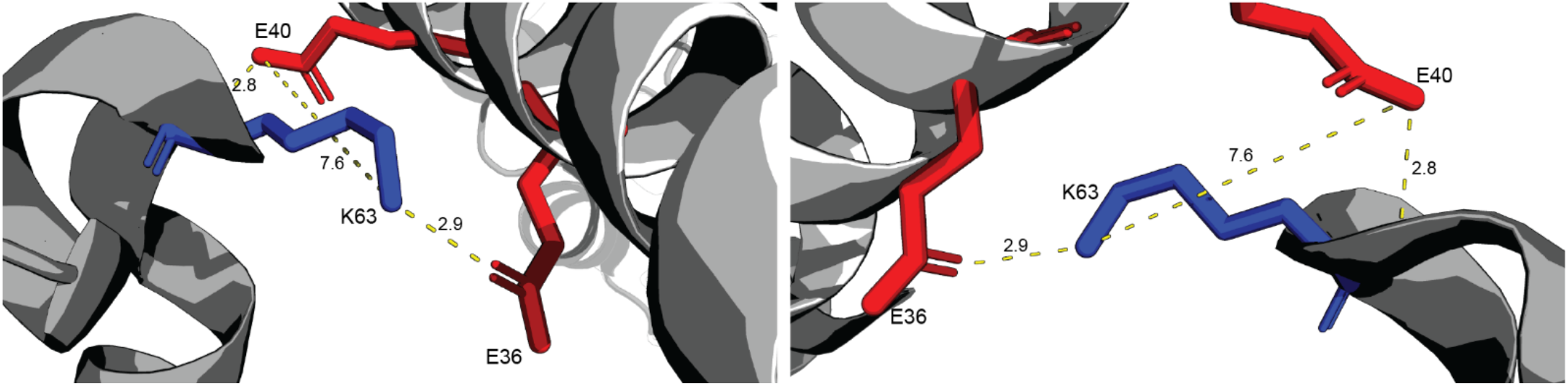
Highlighting structure of conflicting salt bridge between INCENP and Borealin. Highlighting crystal structure between INCENP_1_ and Borealin_2_. Sidechains E36 and E40 of INCENP_1_ have the potential to form a salt bridge with K63 of Borealin_2_. Potential polar contacts and distances between sidechains are labelled.

**Supplemental Figure 6:**
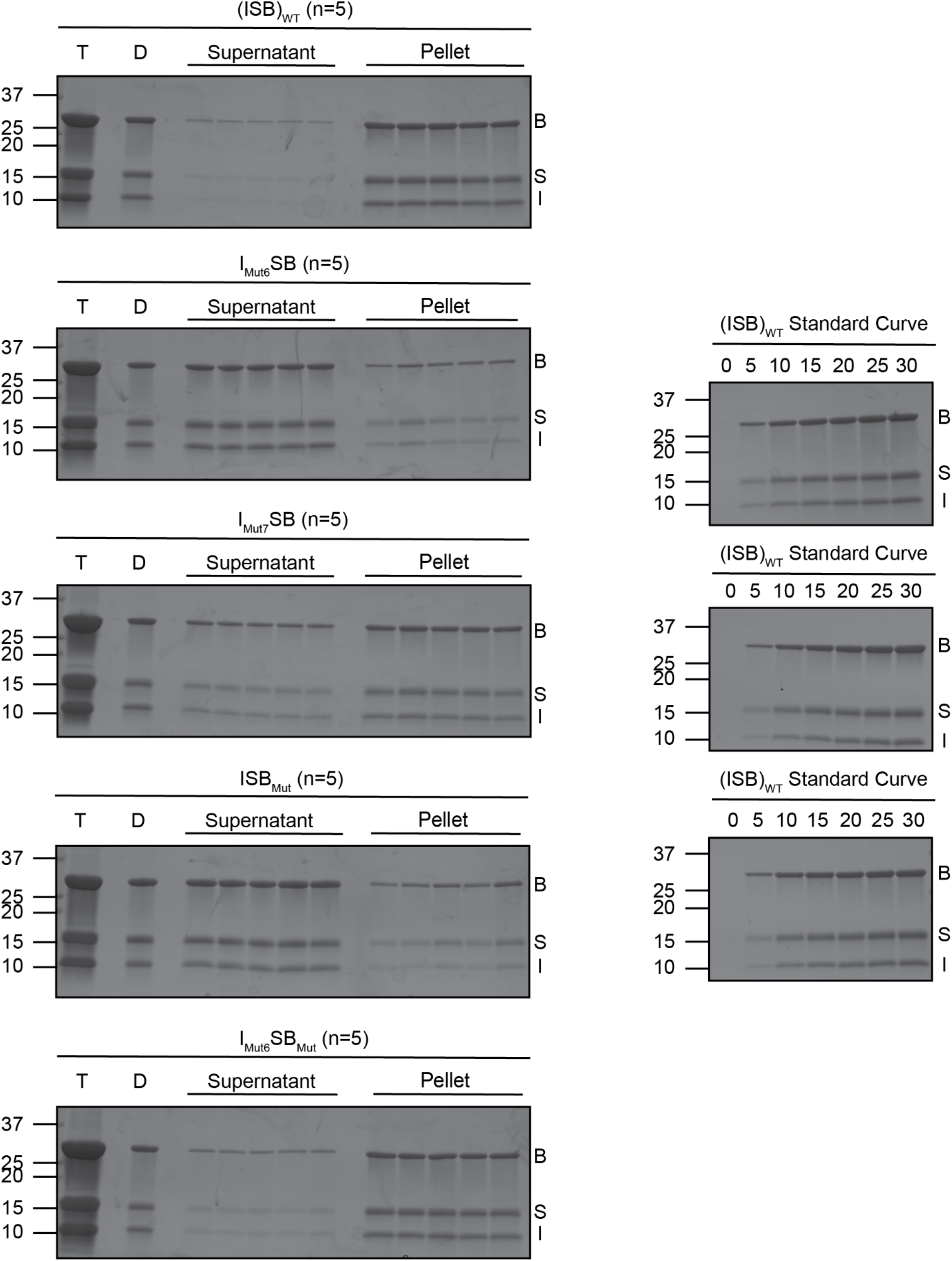
SDS-Gels from Saturation Concentration Experiment. SDS-PAGE gels measuring saturation concentration of (ISB)_WT_, I_Mut6_SB, I_Mut7_SB, ISB_Mut_, and I_Mut6_SB_Mut_ in buffer containing 75 mM NaCl measured using spin-down method. N=5 for all samples. Bands were quantified via GelQuantNET. T = protein after thawing, D = phase separated sample at 25 µM protein and 75 mM NaCl, B = Borealin, S = Survivin, I = INCENP.

**Supplemental Figure 7:**
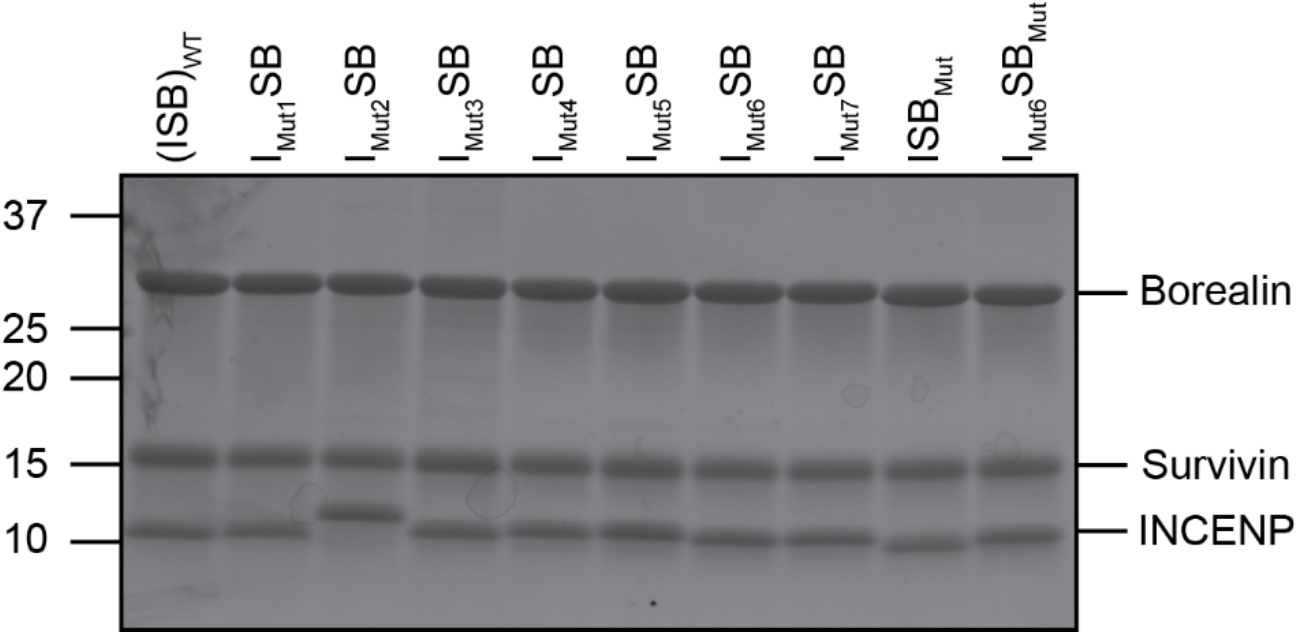
ISB-WT and mutant protein complexes (SDS-PAGE) SDS-Page gel of WT and mutant protein complexes at 1.5 mg/mL.

